# Heterogeneous Sox2 transcriptional dynamics mediate pluripotency maintenance in mESCs in response to LIF signaling perturbations

**DOI:** 10.1101/2025.03.17.643751

**Authors:** Gaochen Jin, Emilia A. Leyes Porello, Jingchao Zhang, Bomyi Lim

## Abstract

The LIF signaling pathway and its regulation of internal factors like Sox2 is crucial for maintaining self-renewal and pluripotency in mESCs. However, the direct impact of LIF signaling on Sox2 transcriptional dynamics at the single-cell level remains elusive. Here, we employ PP7/PCP-mediated live imaging to analyze the transcriptional dynamics of Sox2 under perturbation of the LIF signaling pathway at single-cell resolution. Removal of the LIF ligand or addition of a JAK inhibitor heterogeneously affects the cell population, reducing the number of Sox2-active cells, rather than completely abolishing Sox2 expression. Moreover, Sox2-active cells under LIF perturbation exhibit significant reductions in mRNA production per cell. This reduction is characterized by decreased size and frequency of transcriptional bursting, resulting in shorter duration of Sox2 activity. Notably, cells with reduced or absent Sox2 expression demonstrate a significant loss in pluripotency, indicating that a reduction in Sox2 transcription (rather than a complete loss) is sufficient to trigger the transition from embryonic to an early differentiated state. In LIF-perturbed cells with Sox2 expression reduced to about 50% of non-perturbed levels, we observe a binary behavior, with cells either retaining or losing pluripotency-associated traits. Lastly, we find Sox2 expression is transcriptionally inherited across cell cycles, with Sox2-active mother cells more likely to reactivate Sox2 after mitosis compared to Sox2-inactive cells. This robust transcriptional memory is observed independent of LIF signaling perturbation. Our findings provide new insights into the transcriptional regulation of Sox2, advancing our understanding of the quantitative thresholds of gene expression required for pluripotency maintenance and highlighting the power of single-cell approaches to unravel dynamic regulatory mechanisms.

## Introduction

Mouse embryonic stem cells (mESCs), derived from the inner cell mass of mouse blastocysts, represent a powerful system to investigate mammalian biology across a broad range of fields including gene regulation, disease pathogenesis, and drug development (Evans, 2011; Zakrzewski et al., 2019). The significance and utility of mESCs in both research and clinical applications stem from their key distinct property: pluripotency, the ability to differentiate into any cell types derived from three germ layers (Avior et al., 2016; Kwon et al., 2018). Additionally, their ability to self-renew enables mESCs to divide indefinitely while preserving pluripotency (He et al., 2009). The pluripotent state of mESCs is governed by a highly interconnected pluripotency gene regulatory network (PGRN), centered on Sex-determining region Y-box 2 (Sox2), octamer-binding transcription factor 4 (Oct4), and Nanog (Li and Belmonte, 2017). The level of pluripotent transcription factors (TFs) must be tightly controlled within a narrow range; otherwise, the stability of the cellular state is disrupted, making mESCs prone to differentiation (Festuccia et al., 2013; Li and Belmonte, 2017). For example, cells depleted of Sox2 are likely to exit the pluripotent state and differentiate into trophectoderm-like cells (Masui et al., 2007).

Interestingly, an increase in Sox2 level in KH2 ES cells also results in a shift from an ES-like morphology to a mixed or differentiated one, accompanied by a decrease in the expression of ES cell-associated genes, such as *Nanog*, *Fgf4* and *Utf1*, and an increased expression of genes associated with neuroectodermal, mesodermal and trophectoderm lineages (Kopp et al., 2008). These findings highlight the importance of precise gene regulation in mESCs to maintain pluripotency.

The self-renewal and pluripotency of mESCs are governed by both extracellular factors, like the LIF signaling pathway, and intracellular factors, such as the Sox2 transcription factor (Varzideh et al., 2023). The leukemia inhibitory factor (LIF) signaling pathway contributes to the maintenance of mESC properties by activating multiple pluripotency genes (Ohtsuka et al., 2015; Onishi and Zandstra, 2015; Silva et al., 2008). One such gene regulated by the LIF pathway is Sox2, a key TF essential for sustaining pluripotency and guiding lineage-specific differentiation of ESCs (Schaefer and Lengerke, 2020; Zhang and Cui, 2014). In mESCs, LIF-mediated Sox2 regulation involves three signaling pathways: JAK/STAT3, PI3K/AKT, and SHP2/MAPK pathways (Niwa et al., 2009) (Figure 1D). The JAK/STAT3 and PI3K/AKT pathways affect Sox2 expression by activating Klf4 and Tbx3, respectively. Klf4 directly binds to the Sox2 Control Region (SCR), a Sox2 distal enhancer cluster, to activate Sox2 (Taylor et al., 2022). Tbx3 regulates Sox2 by binding to the regulatory region of Nanog, which in turn targets the SCR (Russell et al., 2015). Conversely, the SHP2/MAPK pathway downregulates Sox2 by suppressing Tbx3 (Hirai et al., 2011; Niwa et al., 2009). These findings highlight the intricate nature of Sox2 regulation through multiple modalities. Yet, the direct effect of LIF pathway modulation on Sox2 transcription remains to be characterized.

**Figure 1.**
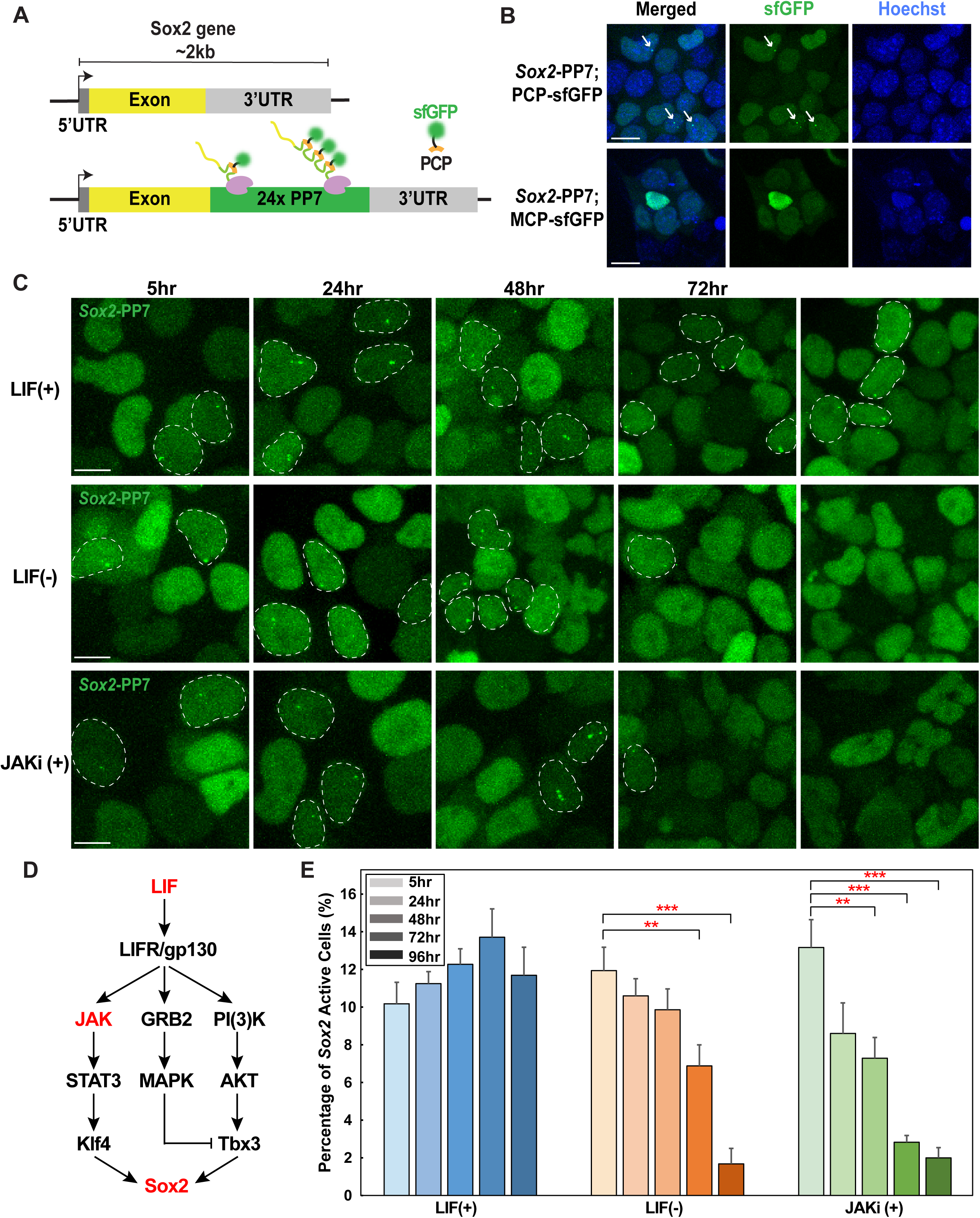
Visualization of Sox2 Nascent Transcripts in Living mESCs Using the PP7 Live Imaging System. (A) Schematic representation of targeting endogenous Sox2 with PP7. 24 copies of the PP7 stem-loop sequence were inserted into the 3’UTR. (B) Snapshots of *Sox2-PP7* tagged E14 cells transfected with PCP-sfGFP (top) and MCP-sfGFP (bottom). Binding between PCP and the PP7 stem loops allows the accumulation of sfGFP, visualized as fluorescent puncta (white arrows). No puncta were observed in MCP-sfGFP transfected cells, due to the absence of binding affinity. Nuclei were stained with the Hoechst dye (blue). (C) Representative snapshots of *Sox2-PP7; PCP-sfGFP* cells without or with LIF-mediated signaling perturbation, denoted as LIF(+) or LIF(-) and JAKi(+) respectively. Cells displaying Sox2 signals are highlighted with white dashed circles. (D) Three LIF-mediated signaling pathways regulating Sox2. In this study, disruption is achieved by removing LIF ligand (red) from the media and adding JAKi to the media (red). (E) Bar chart showing the percentage of Sox2-active cells in each condition, 5-96 hours after each perturbation. The percentage of Sox2-active cells decreases 72 hours after LIF removal (orange) and 48 hours after JAKi addition (green). Sample sizes for total number of cells are as follows: LIF(+) (n = 1376, 2857, 4324, 5373, 2480 for 5-96 hours), LIF(-) (n = 1006, 2040, 4130, 4763, 2439 for 5-96 hours), JAKi(+) (n = 624, 1409, 3234, 4624, 3146 for 5-96 hours), from three independent replicates per condition. Data are presented as mean ± SEM. Statistical significance is indicated as follows: *: P<0.05, **: P<0.01, ***:P<0.001, from the Student’s t-test. See also Figure S1.

Transcription is widely recognized as a stochastic process, producing nascent transcripts as intermittent bursts rather than continuously (McKnight and Miller, 1979; Ross et al., 1994; Thattai and van Oudenaarden, 2004). Such transcriptional bursting has been observed across various organisms and tissue types, including mESCs, along with a significant degree of cell-to-cell heterogeneity in cell populations (Alexander et al., 2019; Leyes Porello et al., 2023; McKnight and Miller, 1979; Ross et al., 1994). However, the range of fluctuations in transcriptional activity that is tolerable to maintain pluripotency or sufficient to induce cell differentiation remains unclear. Moreover, most studies have focused on averaged behaviors across cell populations, overlooking the responses of individual cells (Kopp et al., 2008; Masui et al., 2007; Niwa et al., 2009). Indeed, the regulatory relationship between signaling pathways and downstream pluripotency genes has been widely studied. However, quantitative analysis of how these pathways affect the dynamics of pluripotency gene regulation in individual cells, giving rise to highly heterogeneous expression, is largely lacking (Pascual-Ahuir et al., 2020).

To supplement these population-averaged studies, we employ a live imaging approach to enable single-cell tracking of transcriptional activity in living cells. In this study, we focus on the following key questions: How do individual cells react to perturbation in the LIF signaling pathway? In what way do the transcriptional dynamics of pluripotency genes correlate with the pluripotent state in single cells?

To address these questions, we characterized the transcriptional dynamics of Sox2 under LIF signaling perturbation and examined the impact of changes in Sox2 expression on maintenance of mESC pluripotency. The number of cells expressing Sox2 signals decreased upon LIF pathway perturbation, but there still remained some Sox2-active cells. For cells that still expressed Sox2 under perturbation, mRNA production of Sox2 was significantly reduced in each cell, due to altered transcriptional bursting frequency and size. As a result, cells exhibited a high degree of cell-to-cell heterogeneity in Sox2 levels. A reduction in Sox2 expression was sufficient to disrupt the pluripotency of mESCs, leading to binary cellular behavior, with cells either maintaining or losing pluripotency. Moreover, our live imaging data reveal a transcriptional memory, where a cell with active Sox2 expression has a higher chance of exhibiting active Sox2 transcription in daughter cells after mitosis, regardless of perturbation in LIF signaling pathways. Our work provides a quantitative characterization of how perturbation in the LIF signaling pathway contributes to the heterogeneity in Sox2 transcriptional dynamics that correlate with the pluripotency state in individual cells.

## Results

### PP7 live imaging system allows the visualization of Sox2 nascent transcripts in living mESCs

To examine how the transcriptional activity of Sox2 is affected by the disruption in LIF signaling with a high spatiotemporal resolution, we used the PP7/PCP-mediated live imaging approach that allows visualization of nascent transcripts in living cells. The MS2- and PP7-based live has been widely applied to study transcriptional dynamics across different biological systems, including Drosophila, mouse embryonic stem cells, and human models (Braselmann et al., 2020; Cho et al., 2016; Du et al., 2024; Fukaya et al., 2016). The system functions by incorporating PP7 or MS2 stem-loops into a target gene’s UTRs or introns. During transcription, fluorescently labeled coat proteins (PCP-FP or MCP-FP) bind to these stem-loops, allowing single-cell resolution detection of nascent transcripts in live cells. Using fluorescent in situ hybridization (FISH), previous studies have demonstrated that the insertion of PP7 or MS2 stem-loops minimally perturb endogenous gene transcription, translation, and mRNA stability, while showing a strong correlation between tagged nascent transcripts and endogenous mRNA expression (Braselmann et al., 2020; Das et al., 2018).

Here, we inserted 24 copies of the bacteriophage PP7 sequence into the 3’UTR of the endogenous Sox2 locus using CRISPR/Cas9-mediated genome editing (Figure 1A). The piggybac vector expressing PP7 coat protein (PCP) fused with superfolder GFP (sfGFP) was randomly inserted into the genome of *Sox2-PP7* mESCs. Upon active transcription of *Sox2-PP7*, PP7 sequences form stem loops, where PCP-sfGFP can bind with high affinity, resulting in a bright punctum per allele (Braselmann et al., 2020) (Figure 1B). To ensure that the fluorescent puncta indicate nascent transcripts, a negative control experiment was performed using *Sox2-PP7* mESCs transfected with a plasmid carrying the MS2 coat protein (MCP) fused with sfGFP. As expected, cells transfected with MCP-sfGFP did not exhibit any puncta, indicating that the bright puncta observed in PCP-sfGFP transfected *Sox2-PP7* cells represent Sox2 transcripts (Figure 1B). This assay enabled us to capture nascent transcripts of Sox2 in living mESCs at single-cell resolution.

### Perturbation of the LIF pathway decreases the fraction of Sox2-active cells and increases heterogeneity in cell populations

We first perturbed the LIF signaling pathway by removing the LIF ligand from the culture media. Cells were imaged in a sequential manner at 5, 24, 48, 72, and 96 hours after the removal of LIF ligand. Instead of a complete shutdown of Sox2 transcription, only a fraction of cells was affected upon LIF pathway perturbation. When we quantified the fraction of cells with active Sox2 expression, referred to as Sox2-active cells, a significant decrease was observed (∼42% reduction) 72 hours after removing LIF from the media (Figure 1C, 1E). This reduction became even more pronounced at 96 hours post-LIF removal, showing a five-fold decrease compared to the fraction of transcriptionally active cells 5 hours after LIF removal (Figure 1C, 1E). In contrast, the fraction of Sox2-active cells cultured in LIF media (*Sox2-PP7; PCP-sfGFP*) remained consistent across the five time points (Figure 1C, 1E). A similar trend was observed at a population level (Figure S1).

Among the three characterized LIF-mediated signaling pathways, the activation of JAK/STAT3 pathway alone has been shown to be sufficient to maintain self-renewal in mESCs (Matsuda et al., 1999; Ohtsuka et al., 2015). As such, we analyzed the effects of LIF/JAK/STAT3 signaling pathways on Sox2 transcriptional dynamics (Figure 1D). We cultured the cells in media with 10 µM JAK inhibitor I (JAKi), a potent, cell-permeable, ATP-competitive inhibitor of JAK1 (Thompson et al., 2002). The addition of JAKi yielded results similar to the LIF removal experiments. The fraction of cells expressing Sox2 significantly decreased (∼45% reduction) 48 hours after the addition of JAKi, and this decreasing trend continued progressively over time (Figure 1C, 1E). Notably, there was a time lag of 24 hours between the LIF removal and JAK inhibitor addition methods (Figure 1E). Since JAKi are small molecules that rapidly enter cells and bind the target with high efficiency, this could lead to a faster effect on Sox2 (Hu et al., 2021; Jensen et al., 2023). In addition, LIF ligand, which is previously bound to its receptor before removal, requires time to degrade after its removal, thus delaying the response to this perturbation. Furthermore, the secretion of endogenous LIF by mESCs leads to incomplete removal of LIF from the system, contributing to the slower effect on the LIF removal experiment (Weber et al., 2005). Taken together, both perturbations on the LIF pathway resulted in a decrease in the number of cells expressing Sox2. Rather than homogeneously decreasing Sox2 gene expression in the entire cell population, a gradual and stochastic silencing of Sox2 was observed over time. This variability in response reflects differences in individual cells’ sensitivities to external signals (Figure 1C), thereby contributing to increased heterogeneity within the population.

### Perturbation in LIF pathway leads to reduced mRNA production in Sox2-active cells

In addition to the population heterogeneity following LIF perturbation, single-cell live imaging revealed cell-to-cell variability in transcriptional activity among the Sox2-active cells, regardless of LIF perturbation (Figure 2A, Figure S1B). We investigated whether these Sox2-active cells were resistant to perturbation and maintained the same transcriptional activity as non-perturbed cells or showed changes in transcriptional behaviors. To examine the changes in transcriptional kinetics, we took 5-hour time-lapse videos for each perturbation. We quantified the total mRNA produced in 5 hours for each Sox2-active cell by integrating the transcription trajectories over specific time intervals (Figure 2B). mRNA production is defined as the total integrated fluorescence intensity of Sox2 transcriptional puncta over time, serving as a proxy for cumulative number of nascent transcripts within a given cell (Keller et al., 2020; Pimmett et al., 2021). The mRNA production per cell significantly decreased by approximately 50% at 48 hours after the removal of LIF and 24 hours after the addition of the JAKi, compared to a relatively consistent Sox2 mRNA production in non-perturbed cells (Figure 2D). The observed 24 hours delayed response between the LIF removal and JAKi addition experiments aligns with our previous observations. A previous study using qPCR showed a 50% reduction in Sox2 expression four days after LIF withdrawal, but this averaged data across the population didn’t reveal individual cell behavior (Niwa et al., 2009). Our single-cell analysis addresses this limitation and indicates that perturbation to the LIF-signaling pathway results in not only a decrease in the number of cells expressing Sox2, but also a reduced mRNA production in Sox2-active cells, contributing to the overall decrease in Sox2 expression across the cell population.

**Figure 2.**
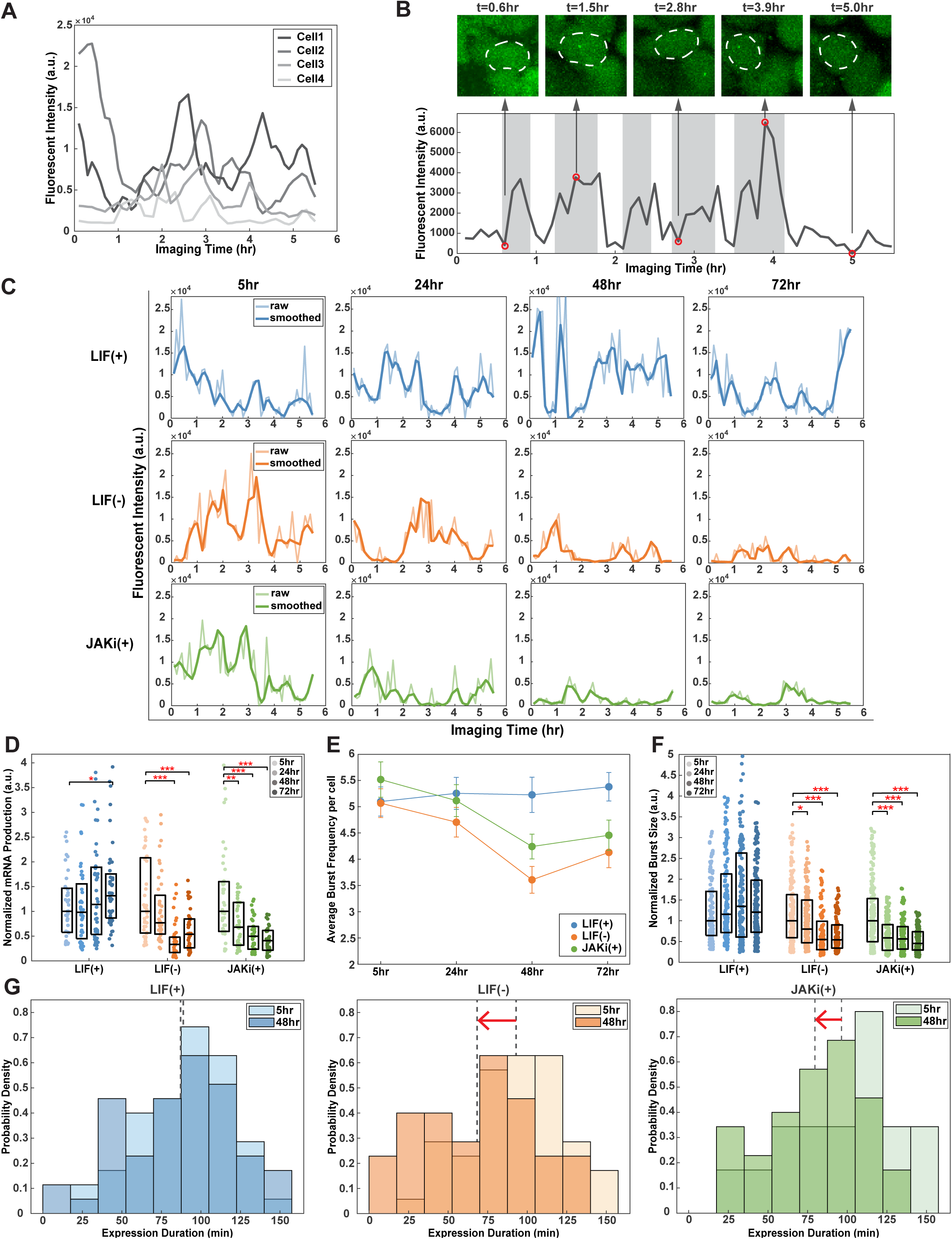
LIF signaling pathway perturbation affects the transcriptional bursting dynamics of Sox2. (A) Transcriptional trajectories of Sox2 vary among Sox2-active cells 72hr after LIF removal, suggesting cell-to-cell variability. Four individual cells’ transcriptional trajectories are shown in different shades of gray. (B) Representative snapshots of a Sox2-active cell 72hr after LIF removal (top) and the corresponding fluorescent intensity profile of the PP7 signal (bottom), displaying stochastic and discontinuous activity over time. The snapshots and the fluorescent intensity trajectory correspond to the same cell. The red circles highlight the representative fluorescent intensities corresponding to the snapshots shown on the top. Each transcriptional burst is highlighted by a gray rectangle. (C) Representative transcriptional trajectories of Sox2-active cells in LIF(+) (top), LIF(-) (middle), and JAKi(+) (bottom) conditions, 5-72 hours after perturbation. Reduced transcriptional bursting is observed following LIF perturbation. The lighter and darker line represents raw and smoothed data, respectively. (D) Boxplots showing mRNA production per cell across different conditions: LIF(+) (n = 61, 60, 62, 66 for 5-72 hours), LIF(-) (n = 60, 60, 56, 54 for 5-72 hours), and JAKi(+) (n = 60, 62, 58, 57 for 5-72 hours). LIF signaling perturbation (LIF(-) and JAKi(+)) result in a significant reduction in mRNA production, with the decreasing trend over time. *: P<0.05, **: P<0.01, ***:P<0.001 from the Student’s t-test. (E) Line plot showing the change in the average burst frequency per cell under each condition, with a significant reduction 48 hours after LIF removal and JAKi addition. Sample sizes for the number of Sox2-active cells are as follows: LIF(+) (n = 61, 60, 62, 66 for 5-72 hours), LIF(-) (n = 60, 60, 56, 54 for 5-72 hours), JAKi(+) (n = 60, 62, 58, 57 for 5-72 hours). Data are presented as mean ± SEM. (F) Boxplots showing normalized burst size across different conditions: LIF(+) (n = 172, 193, 196, 220 for 5-72 hours), LIF(-) (n = 199, 184, 110, 149 for 5-72 hours), JAKi(+) (n = 237, 213, 158, 164 for 5-72 hours). Burst size decreases after perturbation. *: P<0.05, **: P<0.01, ***:P<0.001 from the Student’s t-test. (G) Probability density plots of the duration of active Sox2 expression in each cell 5 hours and 48 hours after perturbation. The duration of Sox2 transcription decreases after perturbation, as indicated by red arrows. The dotted lines demonstrate the average probability density at the two time points. See also Figure S1.

### Reduced Sox2 mRNA level is a result of decreased size and frequency of transcriptional bursting

Given the highly heterogeneous gene expression dynamics observed in individual cells, we next performed a stochastic analysis on individual transcriptional trajectories to investigate the underlying causes of the low mRNA production following disruptions in the LIF signaling pathway. Transcription occurs in a stochastic and discontinuous manner, giving rise to intermittent on and off phases of transcriptional activity, commonly referred to as transcriptional bursting (Leyes Porello et al., 2023; McKnight and Miller, 1979; Ross et al., 1994) (Figure 2B, 2C). Cells exhibited a significantly reduced number of bursts — referred to as bursting frequency—48 and 72 hours post-LIF perturbation, while the number of bursts in non-perturbed cells remained consistent over the same period (Figure 2E). Additionally, both burst size and burst amplitude decreased 48 hours after perturbation (Figure 2F, Figure S1C). The properties of transcriptional bursting are modulated by various factors, including the degree of enhancer-promoter interactions, RNA polymerase II (Pol II) loading rates, TF-DNA interactions, and chromatin modifications (Deng et al., 2022; Leyes Porello et al., 2023). We hypothesized that disruptions in the LIF signaling pathway may influence the regulation of these factors, thereby affecting the bursting properties of Sox2. Consequently, the reduced bursting frequency and size led to a shorter duration of Sox2 activation. Non-perturbed cells maintained a similar distribution of active transcription duration across each 5-hour imaging session, conducted at 5 hours and 48 hours after culture (Figure 2G). However, a noticeable shift toward shorter transcription durations was observed 48 hours after perturbation for both LIF removal and JAKi addition treatments, compared to those observed at 5 hours (Figure 2G). Taken together, our data suggests that perturbation of the LIF signaling pathway affects the transcriptional dynamics of Sox2 by altering burst frequency and size, which leads to a reduced duration of active transcription.

### A significant reduction in Sox2 level leads to binary cellular behavior, with cells either maintaining or losing pluripotency-associated traits under LIF perturbations

Given the integral role of Sox2 in PGRN, we investigated how pluripotency of mESCs is influenced by the LIF-mediated modulation of Sox2. We utilized the live cell antibody of stage-specific embryonic antigen 1 (SSEA1), a cell surface marker of pluripotency, to examine the pluripotent state of mESCs under varied Sox2 expression. In mESCs, a high level of SSEA1 expression is indicative of pluripotency, with differentiation leading to a decrease in SSEA1 levels (Brambrink et al., 2008; Xu et al., 2016). We used the SSEA1 marker conjugated to fluorochrome NL557, which emits red fluorescence, to indicate pluripotency in live mESCs (Figure 3A). Under the non-perturbed condition, more than 90% of cells, regardless of Sox2 activity, showed pluripotency, as indicated by positive SSEA1 signals (Figure S2). However, the percentage of SSEA1-expressing cells decreased by more than 45% 48 hours after the removal of LIF or the addition of JAKi, and by more than 75% 72 hours after treatments (Figure S2). This trend indicates that LIF signaling plays a critical role in sustaining pluripotency, with its perturbation leading to a substantial shift in cell identity.

**Figure 3.**
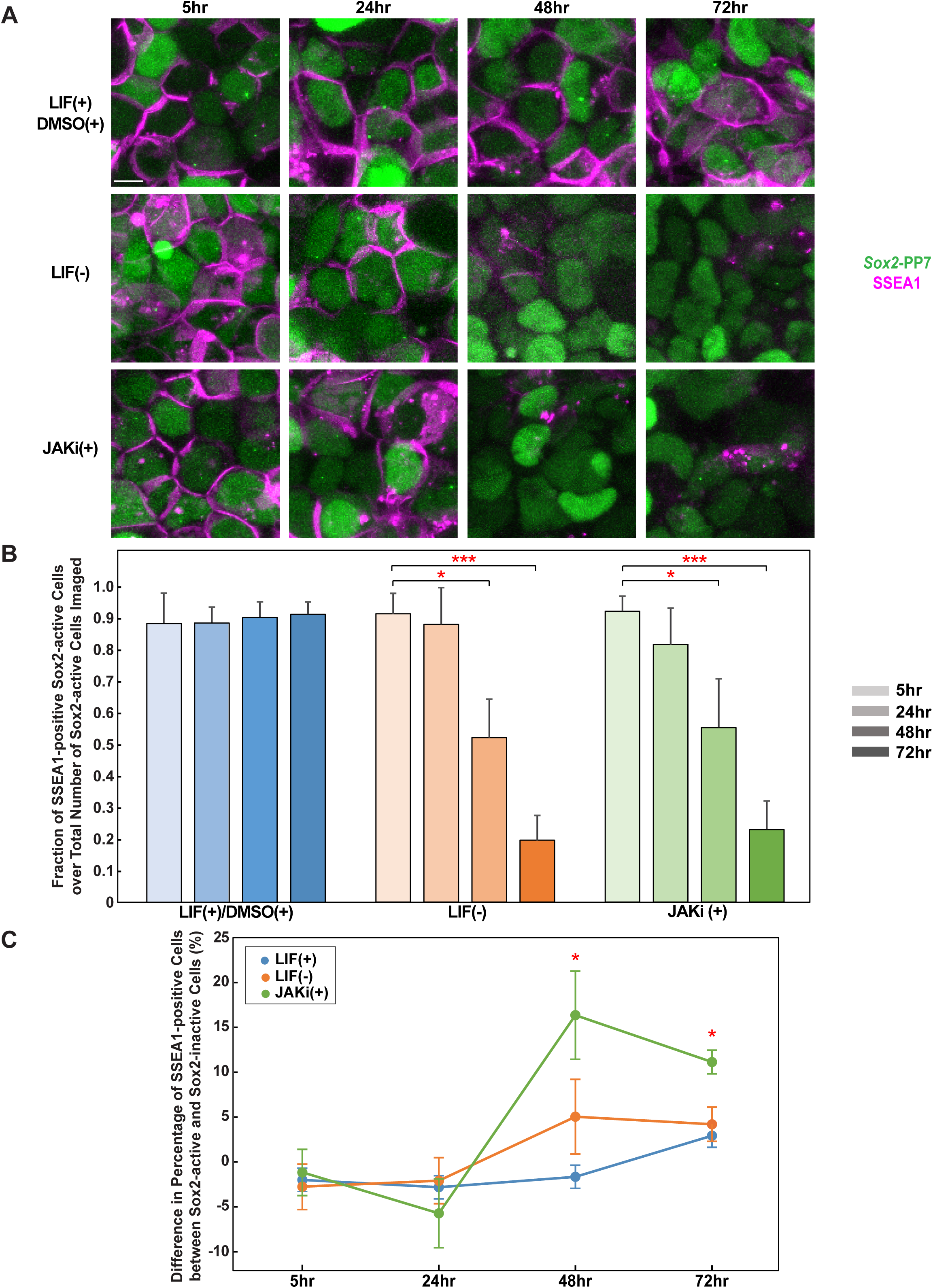
LIF-mediated perturbation results in the loss of pluripotency. (A) Representative snapshots of Sox2-PP7 cells stained with SSEA1, a cell surface marker of pluripotency. A decreased level of SSEA1 expression (magenta) is observed 48 hours after LIF removal and JAKi addition. (B) Bar chart showing the fraction of SSEA1-positive Sox2-active cells relative to the total number Sox2-active of cells in LIF(+)/DMSO(+) (left, n = 285, 208, 190, 159 for 5-72 hours), LIF(-) (middle, n = 237, 215, 108, 106 for 5-72 hours), and JAKi(+) (right, n = 229, 213, 127, 100 for 5-72 hours) conditions, 5-72 hours after perturbation. A significant decrease in the fraction of Sox2-active cells expressing SSEA1 is observed 48 hours after LIF perturbation, indicating the transition of many cells from a pluripotent state to an early differentiate state. Data are presented as mean ± SEM from three independent replicates. *: P<0.05, **: P<0.01, ***:P<0.001 from the Student’s t-test. (C) Line plot showing the difference in the percentage of SSEA1-positive cells between Sox2-active cells and Sox2-inactive cells. The difference became significant 48 hours post-JAKi addition, suggesting that Sox2-active cells are more likely to maintain pluripotency under LIF perturbation. Data are presented as mean ± SEM from three independent replicates. *: P<0.05 from the Student’s t-test. See also Figure S2.

We next analyzed the correlation between Sox2 activity and the pluripotency state. Cells that lost Sox2 expression upon LIF modulation would most likely lose pluripotency. However, would cells that still exhibited Sox2 signals following perturbation, albeit with altered kinetics, retain their pluripotent state? To address this, we analyzed the fraction of SSEA1-positive cells among Sox2-active cells (Figure 3A, 3B). A similar trend was observed, where most cells tended to lose pluripotent traits after perturbation even if they remained Sox2-active (Figure 3A, 3B). We then quantified the difference in the fraction of SSEA1-positive cells between Sox2-active cells and Sox2-inactive cells. This difference became significant 48 hours post-JAKi addition, indicating that cells with Sox2 expression were more likely to remain pluripotent than the cells without Sox2 expression (Figure 3C). Interestingly, the effect of removing LIF was less pronounced compared to adding JAKi (Figure 1D). This less pronounced effect likely arises from the involvement of the JAK/MAPK signaling pathway, which acts as a positive regulator of differentiation and a negative regulator of pluripotency (Hu et al., 2021; Ohtsuka et al., 2015).

The removal of LIF releases the inhibition of Tbx3, a factor known to activate Oct4, potentially resulting in higher Oct4 expression in the absence of LIF compared to the JAKi treatment. It is known that Sox2, Oct4 and Nanog function within an interconnected regulation network, and all play an important role in pluripotency maintenance (Heurtier et al., 2019; Li and Belmonte, 2017; Rodda et al., 2005). Therefore, upregulation of Oct4 could partially compensate for reduced Sox2 levels, helping to sustain pluripotent traits in some cells despite LIF removal.

Building on previous findings that Sox2 stabilizes mESCs in a pluripotent state through Oct4 regulation, our advanced single-cell analysis further underscores the critical role of Sox2 in in maintaining pluripotency and highlights its collaborative function with other TFs in regulating stem cell pluripotency (Masui et al., 2007). Since cells exhibit varying Sox2 expression levels under LIF perturbation, we compared the Sox2 level in cells with or without the pluripotent marker SSEA1. Although Sox2 level was decreased in LIF perturbed cells in general, there was no significant difference in Sox2 intensity between the pluripotent and non-pluripotent cells within the same group (i.e., under the same perturbation) (Figure 2C, S2B). In other words, as long as Sox2 level is dropped to about 50% of the non-perturbed level (Figure 2C), cells can either retain or lose pluripotency-associated traits, exhibiting a binary behavior. In summary, maintaining high levels of Sox2 expression is essential for sustaining the pluripotent state in mESCs. This underscores the complex regulatory mechanisms controlling pluripotency, where Sox2, in concert with other factors, determines the developmental fate of stem cells.

### Transcription memory facilitates Sox2 expression maintenance in mESCs

Previous studies have demonstrated that enhanced Sox2 level leads to a reduction in cell division rates, raising important questions about the stability of Sox2 expression during cell division (Kopp et al., 2008). Specifically, it remains unclear whether Sox2-active cells can maintain Sox2 expression after completing a cell cycle following LIF signaling perturbation. To answer this question, we analyzed the Sox2 transcriptional dynamics before and after mitosis in each condition at each time point from 0 to 72 hours after perturbation. We first sorted the cells into two distinct categories: Sox2-active cells before mitosis and Sox2-inactive cells before mitosis (Figure 4A). Within each group, we examined whether the daughter cells demonstrated any Sox2 expression following mitosis. Among the Sox2-active mother cells, there was a notable tendency for transcriptional reactivation after mitosis, with over 80% of cells displaying Sox2 expression in at least one of the daughter cells (Figure 4B). On the contrary, 90% of Sox2-inactive mother cells did not exhibit any signals following mitosis (Figure 4B). Interestingly, such transcription memory was observed independent of LIF signaling modulations, indicating the inheritance of Sox2 transcription potency despite disruptions in the Sox2 expression level (Figure S3C).

**Figure 4.**
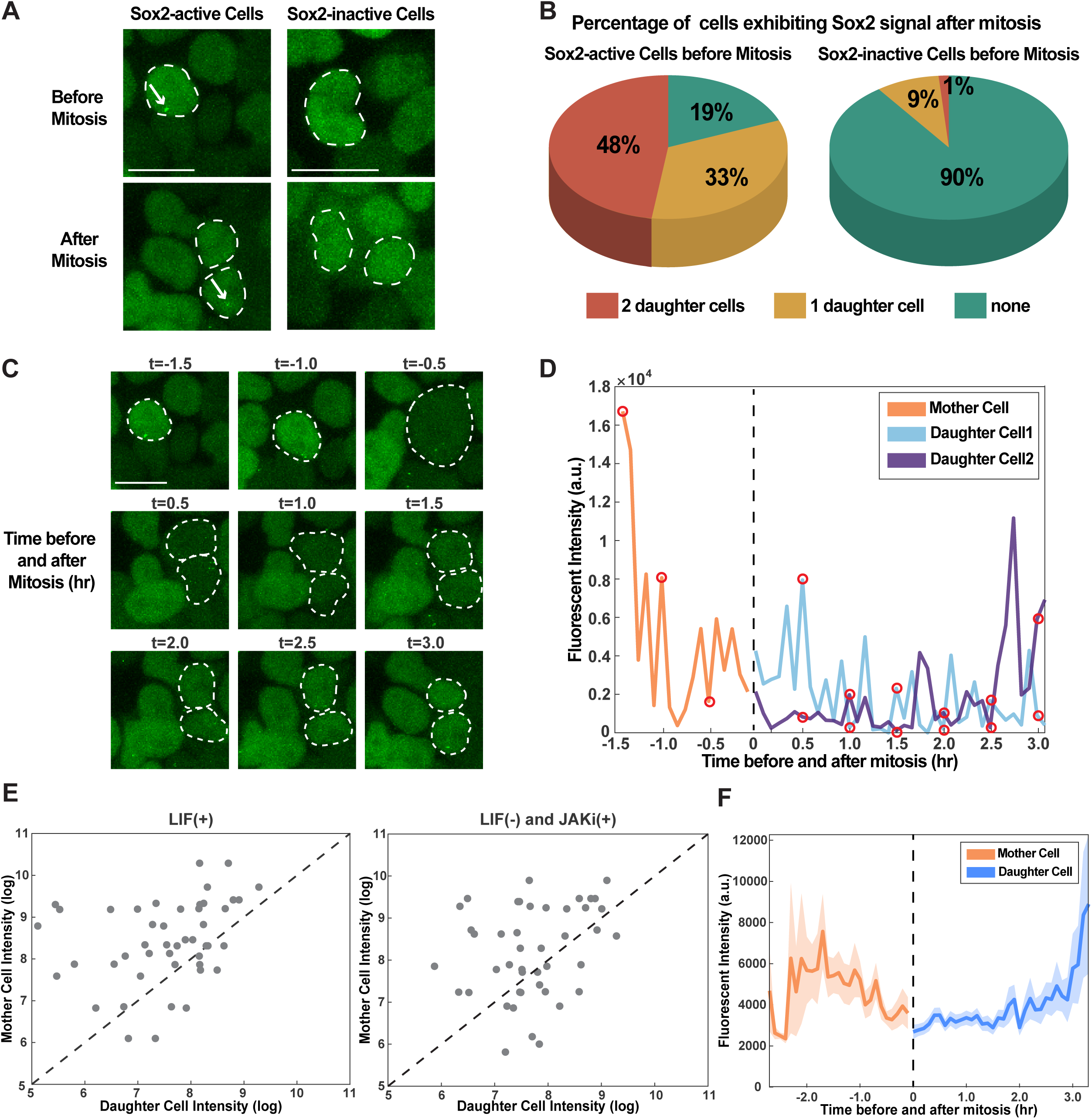
Transcriptional inheritance of Sox2 expression across cell cycles in mESCs. (A) Representative images of non-perturbed *Sox2-PP7* cells (dashed circle) showing active (left) and inactive (right) Sox2 expression, before (top) and after (bottom) mitosis. (B) Pie charts displaying the percentage of cells exhibiting Sox2 signal post-mitosis for Sox2-active (left) and Sox2-inactive (right) cells pre-mitosis in the non-perturbed condition. 48 Sox2-active cells undergoing mitosis and 80 Sox2-inactive cells undergoing mitosis were analyzed. Over 80% of Sox2-active pre-mitosis cells exhibit Sox2 in at least one daughter cell, while 90% of Sox2-inactive cells do not show transcription after mitosis. (C) Snapshot of a cell undergoing mitosis, highlighted by dashed circles. (D) Corresponding transcriptional trajectories of the mother cell (orange) and two daughter cells (cyan and purple). Red circles represent the fluorescent intensities of *Sox2-PP7* corresponding to the snapshots in (C). The dotted line shows the onset of mitosis. (E) Log-log scatter plot of the Sox2 activity between mother and daughter cells for LIF(+) cells (left, n = 23) and cells with LIF perturbation (right, n =25). The transcriptional activity of a mother cell is generally higher than that of the daughter cells. (F) Average transcriptional activity of Sox2 in mother cells (n = 38) (orange) and daughter cells (n = 76) (blue), with a dotted line indicating the onset of mitosis. Shaded area represents SEM. Daughter cells show lower Sox2 expression during mitosis exit, followed by an increase in fluorescent intensity thereafter. See also Figure S3.

To further characterize the observed transcription memory, we tracked the Sox2 transcription in mother cells and corresponding daughter cells (Figure 4C, 4D). Both mother cells and daughter cells exhibited Sox2 transcription in a stochastic and discontinuous manner regardless of LIF perturbation, indicating that transcriptional bursting occurs in both short (within a few hours) and long (spanning cell cycles) time scales (Phillips et al., 2019) (Figure 4D, S3). We found that daughter cells were likely to exhibit lower Sox2 expression compared to their mother cells both in non-perturbed and perturbed (LIF-removal and JAKi-addition) conditions (Figure 4E). This could be explained by the fact that during mitotic exit, Sox2 expression stays at a low level (Palozola et al., 2017a) (Figure 4F). Our data demonstrate that Sox2 expression can be transcriptionally inherited from mother cells, regardless of the LIF signaling pathway perturbation. We note that while Sox2 transcriptional inheritance does not rely on LIF signaling pathway, both the overall level and stability of Sox2 expression are still significantly influenced by LIF. This indicates that LIF plays an important role in maintaining Sox2 expression, but the fundamental mechanism of Sox2 transcriptional inheritance across cell cycles remains functional even when LIF signaling is perturbed.

## Discussion

The LIF signaling pathway plays a crucial role in maintaining pluripotency in mESCs, acting through both direct and indirect regulatory mechanisms (Ohtsuka et al., 2015; Onishi and Zandstra, 2015; Silva et al., 2008; Varzideh et al., 2023). While some transcription factors are directly controlled by JAK/STAT signaling, such as Klf4, their roles in pluripotency can be functionally redundant due to overlapping regulatory networks (Jiang et al., 2008; Yamane et al., 2018). In contrast, Sox2 is an essential member of the core pluripotency transcription factor network, working synergistically with Oct4 and Nanog to maintain the pluripotent state (Li and Belmonte, 2017; Masui et al., 2007). Additionally, Sox2 expression is regulated through multiple LIF-mediated signaling pathways via Klf4, Tbx3 and other factors, integrating LIF signals into the broader pluripotency network (Niwa et al., 2009). Hence, monitoring Sox2 activity allows us to capture the integrated effects of LIF signaling on pluripotent network.

Beyond its role in the LIF response, Sox2 is one of the three core transcription factors governing the pluripotency gene regulatory network (PGRN) and is essential for both the preservation of pluripotency and the orchestration of cell fate decisions during embryogenesis (Schaefer and Lengerke, 2020; Varzideh et al., 2023; Zhang and Cui, 2014). It has been established that transcription of many genes, including Sox2, occurs in bursts, and such stochastic transcriptional dynamics have profound implications for the regulation of pluripotency in mESCs (Alexander et al., 2019; Leyes Porello et al., 2023). Yet, much remains to be understood about the extent of transcriptional fluctuations that can be tolerated to maintain pluripotency or the range that induces cell differentiation. In this study, we use the PCP/PP7-based single-cell live imaging to elucidate how modulations of the LIF signaling pathway affect downstream Sox2 transcriptional dynamics, and the subsequent impact on pluripotency maintenance.

LIF signaling perturbation alters Sox2 transcriptional dynamics by decreasing the frequency and size of transcriptional bursts, indicating a regulatory effect of the LIF signaling pathway on pluripotency gene expression dynamics. The relationship between LIF signaling and transcriptional bursting suggests that LIF interacts with the transcriptional machinery and its regulatory elements, such as promoters and enhancers, to sustain the high transcriptional output necessary for maintaining pluripotency. For example, the SCR, a strong distal enhancer of Sox2, contains several TF binding sites, including two Klf4 sites (Taylor et al., 2022). Given that Klf4 is a direct target of the LIF/STAT3 signaling pathway and is solely influenced by this pathway, further research is needed to explore the correlation between Klf4 protein binding and Sox2 transcription. Combining our live imaging technique with protein tagging approaches would offer a more comprehensive understanding of how pluripotency genes, along with other factors—including extrinsic signaling pathways and intrinsic regulatory elements (e.g., promoters and enhancers)—orchestrate the transcriptional dynamics and the maintenance of pluripotency. Furthermore, while we have used a simple two-state model to characterize transcriptional bursting, it is sensitive to noise and often fails to capture some critical factors influencing complex gene expression in the mammalian system (Cao et al., 2020; Jia and Grima, 2023). To better account for these complexities, the two-state model has been modified into variations like the delayed telegraph model, three-state model, and mechanistic multi-state model (Braichenko et al., 2021). It would be of interest to see if these advanced models fit our Sox2 expression data better and provide additional insights.

Our findings reveal that LIF perturbation leads to a significant decrease in the fraction of cells expressing the pluripotent marker SSEA1, indicating a potential reduction in pluripotency-associated traits. Interestingly, cells expressing Sox2 are more likely to maintain pluripotency-associated traits under LIF perturbation, emphasizing the heterogeneous sensitivity to LIF perturbation. While Sox2 is essential for pluripotency, the intensity of Sox2 level is not significantly different between SSEA1-positive vs. SSEA1-negative cells — both exhibiting lower Sox2 level compared to non-perturbed cells. Sox2 transcriptional regulation at low levels seems to be governed by discrete ’on’ or ’off’ states, contributing to a more deterministic control mechanism, rather than a continuous gradient of expression. These results demonstrate that when Sox2 remains at a basal level, it functions in a binary manner to retain or lose pluripotency-associated traits. By leveraging single-cell analysis, we provide a detailed perspective on the transcriptional dynamics of Sox2, providing a gene-level understanding of the regulatory mechanisms that underlie pluripotency and cell fate decisions.

We note that perturbations in the LIF pathway affect multiple genes beyond Sox2, including Klf4, Oct4, and Nanog (Li and Belmonte, 2017; Niwa et al., 2009). This suggests that the observed effects on pluripotency may result from the combined impact of altered transcriptional dynamics across several genes. Indeed, our results show that the effect on pluripotency was less pronounced for the LIF removal experiment compared to the more specific JAKi addition experiment, possibly due to the indirect activation of Oct4 upon LIF removal. Additional studies using approaches like single-cell RNA-seq or multiplexed monitoring of several genes will be necessary to uncover the combinatorial contributions of pluripotency genes on pluripotency maintenance.

We also reveal that Sox2 expression can be transcriptionally inherited across cell cycles, independently of LIF signaling status. A previous study using a mathematical model demonstrated that the intrinsic heterogeneity in gene expression is caused by multi-generational memory, where a mother cell and its two daughter cells have comparable mean transcriptional activities (Phillips et al., 2019). However, we demonstrate that daughter cells generally show lower expression levels than mother cells, regardless of LIF signaling perturbation. A recent study showed that many genes are expressed at a low level during mitosis, rather than being silenced (Palozola et al., 2017a). While enhancer accessibility is lost during mitosis, promoter accessibility is largely maintained, allowing the promoter and its target gene to form mitotic expression units that preserve transcription patterns at a low level. During mitotic exit, enhancers are reactivated, and the long-range enhancer-promoter interactions gradually resume, resulting in gene expressions that remain low initially but increase during the transition from mitosis to interphase (Ito and Zaret, 2022; Palozola et al., 2017b). Consistent with previous findings, our results specifically show that Sox2 expression follows this pattern, remaining low during mitosis followed by a gradual increase in signal. Notably, we find that this transcriptional pattern is independent of LIF signaling perturbations. Therefore, we propose that while LIF plays an important role in maintaining Sox2 expression, the fundamental mechanism of Sox2 transcriptional inheritance across cell cycles remains functional even when LIF signaling is perturbed, reflecting the stable intrinsic properties of promoters.

Taken together, our research underscores the significant impact of LIF signaling perturbation on the downstream Sox2 transcriptional dynamics and the maintenance of pluripotency in mESCs. Utilizing live-cell imaging techniques that allow single-cell resolution, we reveal significant heterogeneity in Sox2 expression following disruptions in the LIF pathway. Our results indicate that Sox2 bursting dynamics, modulated by signaling pathways like the LIF pathway, play a critical role in pluripotency maintenance. Further investigations incorporating multiplexed gene analysis and more sophisticated transcriptional models will be essential to fully elucidate the complex interactions among pluripotency regulators and their influence on cellular development. Overall, our findings emphasize the vital role of transcriptional regulation in mESC pluripotency, setting the stage for the development of groundbreaking models and therapeutic approaches that can precisely manipulate stem cell fate, ultimately propelling advancements in regenerative medicine.

## Methods

### Cell culture and treatments

All mESC lines were cultured in LIF/serum medium supplemented with DMEM (Gibco, 11995065), 10% Fetal bovine serum (Hyclone, SH30071.03), 1 mM MEM nonessential amino acids (Invitrogen, 11140035), 0.1 mM β-mercaptoethanol (Gibco, 21985023), 1000U/ml LIF(Millipore, ESG1107) and 100U/ml Penicillin-Streptomycin (Gibco, 15140122) on 0.1% gelatin-coated plates in a humidified incubator at 37°C and 5% CO_2_. To perturb LIF-related signaling pathways, mESCs were either treated with 10 μM JAK inhibitor I (Carbiochem, 457081037) or with LIF-depleted medium.

### Cell line generation

The targeting vectors were generated that incorporated a 24x PP7 sequence and a puromycin selection cassette, flanked by the homology arm sequences corresponding to the ∼500bp upstream and downstream of the guide RNA target site. The DTA negative selection sequence was inserted downstream of the right homology arm. gRNAs were designed using CRISPick with Mouse GRCm38, CRISPRko and SpyoCas9 (cite). The PX458 plasmid backbone was used to insert the oligos containing the gRNA sequence. Both the donor and the gRNA-containing vectors were purified using the NucleoBond Xtra midiprep kit (Takara, 740410.50) for further transfection. The ePB-Ubc-PCP-sfGFP-Hygro plasmid was generated by substituting the MCP sequence to the PCP sequence in the ePB-Ubc-MCP-sfGFP-HygroR plasmid, kindly provided by Dr. Jingchao Zhang from the Zaret Lab at Penn Medicine.

Wildtype E14 cells were transfected with 2 μg of the gRNA vector and 1 μg of the donor vector using opti-MEM (Gibco, 31985070) and Lipofectamine 3000 Transfection Reagent (ThermoFisher, L3000001). After two days, transfected cells were subjected to antibody selection for a duration ranging from 7 to 14 days depending on the specific antibody used. Cells that went through antibody selection were replated at a diluted concentration to pick individual colonies. Genomic DNA was extracted from each colony and genotyped to validate the correct insertion of the target sequence in the designated location. Promising clones with bi-allelic integrations were co-transfected with the ePB-Ubc-PCP-sfGFP-HygroR plasmid and 1μg PiggyBac enzyme PBase using opti-MEM (Gibco, 31985070) and Lipofectamine 3000. Two days following transfection, cells were treated with 100 μg/ml Hygromycin B (Corning, 30240CR) for 14 days. Individual colonies were picked after hygromycin selection and screened under a confocal microscope. The colony exhibiting the highest signal-to-noise ratio of PP7 signal compared to the PCP background level was chosen for further experiments.

### Live imaging

Cells were plated onto a 12-Well Glass Bottom Plate (Cellvis, P121.5HN) coated with Matrigel (Corning, 354277) either 2 or 3 days before imaging with desired concentrations. The cells were cultured at 37°C and 5% CO_2_ during the imaging process using a stage incubator. All images were taken using a Zeiss LSM800 confocal laser scanning microscope and Plan-Apochromat 40x/1.3 numerical aperture oil-immersion objective. Nascent transcripts were visualized using a 488 nm laser. For calculating the fraction of cells with Sox2 signal and the pluripotency test, snapshots were taken at six random positions for three experimental replicates. For transcriptional activity analysis, images were taken every 5 minutes for 5 hours at six random positions. Maximum projection of 40 z-stacks with 0.60 μm per stack was used for further analysis. Each single 512 x 512-pixel image was captured in 16-bit. The same laser settings were maintained throughout all experiments. For perturbation related to LIF-signaling, cells were either treated with 10 μM JAK inhibitor I (Carbiochem, 457081037) or with LIF-depleted medium. The time when perturbation was induced was defined as time 0 and the cells were imaged at specific time points in a sequential manner (5 hours, 24 hours, 48 hours and 72 hours after the treatments). Each time-lapse movie spanned a duration of 5 hours.

To visualize the pluripotency state of mESCs in living cells, GIoLIVE mouse live cell antibody targeting the SSEA1 was used (Bio-Techne, NLLC2155R). SSEA1 antibody was conjugated to fluorochrome NL557. The antibody was diluted 1:50 to a final concentration of 1X in cell culture media. A staining volume of 2mL was added to one well in a 12-well plate. After 30 minutes of incubation, the staining media was replaced with fresh media without antibodies. Then cells were used directly for imaging. Pluripotent stem cells exhibit the fluorescent signal of SSEA1 while early differentiated cells do not fluoresce.

### Image analysis

All image processing methods and analyses were conducted using MATLAB (R2019b, MathWorks) and Fiji ImageJ. Fluorescence histograms adjustments were applied only for visualization, while all quantitative analyses were conducted using raw intensity values.

The images requiring analysis were concatenated and subjected to maximum projection using Fiji ImageJ. Maximum intensity projection images were utilized for cell segmentation and the tracking of nascent transcript intensities. The code for cell segmentation and signal tracking was developed using MATLAB. Object segmentation was performed on each frame of the time series and produced centroid indices and pixel lists associated with each object (cell) in the image. To track objects across time frames and correctly assign cell identities over the time course, a key criterion was applied: the object areas between the two consecutive frames must overlap by >60%. Once segmented objects were assigned proper cell identities across frames, the MS2 signals were extracted in each cell and frame and tracked over time.

### MS2 signal identification

The MS2 signal identification and tracking was performed following a similar algorithm as cell segmentation but instead of applying an area overlap criterion, signal identity was assigned based on a centroid distance threshold. A scalar threshold of 7 pixels was applied such that signals in consecutive frames with centroid distances < 7, were assigned the same identity. For signal quantification, to account for background variation, we subtracted the mean fluorescence intensity in the nucleus from each detected punctum. The puncta that persisted for more than two consecutive frames were considered as valid transcriptional events. To further validate our quantification method, we conducted manual curation of signal tracking and confirmed the consistency of our analysis.

### Data smoothing and transcriptional activity calculation

Following the cell identification and signal quantification, the trajectory data were smoothed using the “smoothdata()” function by “rlowess” method over intervals of 5 frames. Subsequently, a threshold intensity value (A.U.) was applied to facilitate the identification of active transcription events. Cells were sorted as Sox2-active or Sox2-inactive cells based on the presence of transcriptional puncta during the imaging window.

### Quantification of mRNA production and bursting activity

For each individual cell, mRNA production was quantified by integrating the area under the transcriptional trajectory using the trapezoid rule. To facilitate comparisons within each condition (LIF(+), LIF(-), and JAKi(+)), mRNA production values were normalized to the mRNA production at 5 hours in each condition. To quantify transcriptional bursting kinetics in each Sox2-active cell, bursts were identified using local maxima and minima of the transcriptional trajectory. A burst was defined when the fluorescent intensity difference between a local maximum and its two adjacent minima exceeded a defined threshold. This threshold was empirically determined to clearly distinguish between active (“on”) and inactive (“off”) transcription states. Bursting frequency was calculated as the number of detected bursts per cell within the imaging period.

The average bursting frequency for each condition was determined by averaging the number of bursts per cell. Burst size was quantified as the integrated area under the transcription trajectory for each burst, and burst amplitude was measured as the maximum fluorescence intensity of the burst. Both burst size and amplitude were normalized to the value for the 5-hour time point in each condition. The duration of Sox2-active expression was quantified as the time during which the fluorescence intensity exceeded the threshold for each cell. The results are presented as a probability density plot.

## Statistical analysis

Statistical analysis was performed using MATLAB and Excel. An unpaired two-tailed Student’s t test was used to determine statistical significance when comparing two independent groups. Unless otherwise specified, data represents the mean ± standard error of the mean (SEM) from three independent experiments. Each experiment was performed in triplicate unless stated otherwise.

## Resource Availability

### Lead contact

Further information and requests for resources and reagents should be directed to and will be fulfilled by the lead contact, Bomyi Lim.

### Materials availability

Plasmids generated in this study are available upon request.

### Data and code availability

The code used in this study is available upon request.

## Acknowledgments

We thank members of the Lim lab for helpful discussions and comments on the manuscript. E14 embryonic stem cells were kindly provided by the Zaret Lab at the Perelman School of Medicine, University of Pennsylvania. J.Z. was supported by a Human Frontiers Science Program fellowship (LT000761/2019-L). E.A.L.P. is partially funded through the University of Pennsylvania Fontaine Society. This study is funded by NIH R35GM133425 awarded to B.L.

## Author contributions

G.J. and B.L. designed the research and performed the research; E.A.L.P wrote Matlab code for data analysis; J.Z. mentored G.J. for mammalian cell culture and generated plasmids; G.J. analyzed data; G.J. and B.L wrote the initial draft of the manuscript; G.J., E.A.L.P, J.Z., and B.L. reviewed and edited the manuscript.

## Declaration of interests

The authors declare no competing interests.

---

**Figure S1. Visualization of Sox2 Nascent Transcripts in Living mESCs Using the PP7 Live Imaging System.** (A) Population-scale visualization of *Sox2-PP7; PCP-sfGFP* cells without or with LIF-mediated signaling perturbation, denoted as LIF(+) or LIF(-) and JAKi(+) respectively. (A) (B) Transcriptional trajectories of Sox2-active cells under non-perturbed conditions, suggesting cell-to-cell variability. Four individual cells’ transcriptional trajectories are shown in different shades of gray. Variability in Sox2 transcriptional activity was observed among Sox2-active cells, regardless of LIF perturbation. (C) Boxplots showing normalized burst amplitude across different conditions: LIF(+) (n = 172, 193, 196, 220 for 5-72 hours), LIF(-) (n = 199, 184, 110, 149 for 5-72 hours), JAKi(+) (n = 237, 213, 158, 164 for 5-72 hours). Burst amplitude decreases following perturbation. *: P<0.05, **: P<0.01, ***:P<0.001 from the Student’s t-test.

**Figure S2. LIF-mediated perturbation results in the loss of pluripotency.** (A) Representative snapshots of *Sox2-PP7* cells stained with SSEA1 under non-treated (top) and DMSO-treated (bottom) conditions. No significant differences were observed between the two groups, confirming that DMSO serves as an appropriate control for the experiment. (B) Bar chart showing the fluorescence intensity of Sox2 in cells with SSEA1 (magenta) and cells without SSEA1 (gray) in LIF(+)/DMSO(+) (left), LIF(-) (middle), and JAKi(+) (right) conditions, 5-72 hours after perturbation. No significant differences in Sox2 intensity were observed between pluripotent and non-pluripotent cells across all conditions. Sample sizes are as follows: LIF(+)/DMSO(+) (total number of Sox2-active cells with SSEA1 = 287, 196, 187, 151 for 5-72 hours, total number of Sox2-active cells without SSEA1 = 21, 21, 17, 11 for 5-72 hours), LIF(-) (total number of Sox2-active cells with SSEA1 = for 235, 203, 67, 26 for 5-72 hours, total number of Sox2-active cells without SSEA1 = 16, 26, 49, 83 for 5-72 hours), JAKi(+) (total number of Sox2-active cells with SSEA1 = for 226, 190, 84, 30 for 5-72 hours, total number of Sox2-active cells without SSEA1 = 16, 39, 59, 80 for 5-72 hours). Data are presented as mean ± SEM.

**Figure S3. Transcriptional inheritance of Sox2 expression across cell cycles in mESCs.** (A) Snapshot of a cell 48 hours after JAKi addition undergoing mitosis, highlighted by dashed circles. (B) Corresponding transcriptional trajectories of the mother cell (orange) and two daughter cells (cyan and purple). Red circles represent the fluorescent intensities of Sox2-PP7 corresponding to the snapshots in (A). The dotted line shows the onset of mitosis. (C) Pie charts showing the percentage of cells with Sox2 signals post-mitosis for Sox2-active (top) and Sox2-inactive (bottom) cells pre-mitosis, under non-perturbed (left) and perturbed (right) conditions (n = 23 number of non-perturbed Sox2-active cells undergoing mitosis; n = 25 perturbed number of Sox2-active cells undergoing mitosis; n = 43 number of non-perturbed Sox2-inactive cells undergoing mitosis; n = 37 perturbed number of Sox2-inactive cells undergoing mitosis). No significant differences observed between the perturbed and non-perturbed groups, suggesting Sox2 transcriptional inheritance is independent of LIF perturbations. (D) Log-log scatter plots comparing Sox2 activity between mother and daughter cells, shown for 5-24 hours after LIF perturbation (left) and 48-72 hours after LIF perturbation (right) (n = 26 number of cells undergoing mitosis after 5-24 hours after perturbation; n = 22 number of cells undergoing mitosis after 48-72 hours after perturbation). Sox2 transcriptional activity is consistently higher in mother cells than in their daughter cells, regardless of the duration of LIF perturbation.

## Notes

### Competing Interest Statement

The authors have declared no competing interest.

